# Context-aware Multi-Property Antibody Predictor: a Novel Framework Integrating Text and Protein Language Models

**DOI:** 10.64898/2026.01.07.698270

**Authors:** Luca Giancardo, Melih Yilmaz, Edward Lee, Ke Ren, Yue Zhao, Gordon Trang, Kemal Sonmez, Lan Guo, Nina Cheng

## Abstract

Recent advances in Machine Learning have transformed antibody development through in-silico models, accelerating therapeutic candidate identification. However, challenges persist: rapid adaptation of property predictors to laboratory-specific assays with incomplete datasets; batch effects introducing systematic bias; assay costs necessitating efficient unseen property prediction.

We introduce a novel multi-modal architecture featuring specialized tokenization and embedding projection that integrates text and protein language models (pLM) and a learning strategy to enable in-context learning for multi-property prediction without learning shortcuts. Our framework enables prompting without dictionary merging across modalities, creating a compact model capable of in-context learning for multi-property prediction. The orchestrating model avoids pLM-to-text projection while enabling inference-time adaptation without retraining.

Using 876,898 antibodies with batch effect simulation, our architecture achieved Spearman’s ρ>0.8 across multiple developability properties, significantly outperforming fine-tuned multimodal-LLMs and showed the ability of leveraging correlation between properties for prediction. This approach has the potential to address critical antibody development challenges.

## Introduction

Antibody therapeutics have revolutionized modern medicine, representing the fastest-growing class of therapeutic proteins with applications spanning oncology, autoimmune diseases, and infectious diseases.^1^ However, the traditional antibody development process remains highly time-consuming and resource-intensive, often requiring years of optimization through iterative rounds of experimental screening and affinity maturation.^2^ The complexity of this process has intensified the need for computational approaches that can accelerate antibody discovery while reducing experimental costs and development timelines. One of the critical aspects for a successful antibody campaign is the developability assessment of antibodies, this involves properties such as stability, solubility, and aggregation propensity. ^3–5^

Historically, assessing developability has relied on time-consuming laboratory-based assays. As a result, significant resources have been dedicated to creating computationally based tools and machine learning models that can estimate or forecast developability parameters using antibody sequence and structural data, providing more practical alternatives to experimental methods. ^6–11^ Recent advances in machine learning have fundamentally transformed the landscape of antibody development through in-silico models that can predict and optimize antibody properties and analysis of large-scale antibody repertoire data, facilitating the identification of patterns and signatures associated with specific properties. ^12,13^ This has been further boosted by the emergence of protein language models (pLMs), offering enhanced capabilities for general purpose protein representation ^14–16^ and antibody-specific representations. ^17–21^ These models, trained on vast protein sequence databases, have demonstrated remarkable performance in capturing evolutionary relationships and functional patterns within protein families.

Multi-task and multi-modal architectures, also able to integrate text, have emerged as particularly promising approaches for biological applications, as they can simultaneously leverage multiple sources of information to provide more comprehensive and robust predictions particularly for chemical reactions for small molecules and proteins. ^22–25^ These multi-modal architectures typically require like some type of fine-tuned specialized head or full fine tuning to a single task (e.g. with BioT5^23^), this hinders the ability to learn from relationships between multiple properties for prediction. Studies with direct multi-task ability have only limited or no validation on antibody property data (e.g. Regression Transformer^25^ or protLLM^22^). A notable advancement is TxGemma, ^24^ a specialized multi-modal large language model that demonstrates excellent performance across therapeutic development tasks including antibodies and allowing for changing prediction tasks by changing input text prompts, thereby removing the need of specialized single-task fine-tuning.

Despite these significant advances, three critical challenges continue to limit the broader adoption and effectiveness of current in-silico antibody development approaches. First, there is an urgent need for rapid adaptation of property predictors to laboratory-specific assays and mutation campaigns, particularly when working with incomplete or limited datasets that are common in early-stage therapeutic development.^26^ In-context learning has shown promise for addressing some of these challenges by enabling models to adapt to new tasks and datasets without extensive retraining. While ProtLLM does suggest the potential for encoding multiple proteins and properties in its context for n-shot learning at inference time, the application of in-context learning principles to multimodal biological data, particularly in the context of antibody property prediction with explicit batch effect handling, remains largely unexplored. Such a model would allow for rapid adaptation to new laboratory conditions.

Second, batch effects in laboratory data introduce systematic biases that can severely compromise model performance and generalizability across different experimental conditions, instruments, and laboratories. ^27^ While numerous methods have been developed for correcting batch effects in various biological assays,^26–28^ their integration with modern deep learning architectures, particularly in the context of multimodal protein language models, has not been addressed. This integration is essential for developing robust models that can operate effectively across diverse experimental conditions without extra processing steps and extensive retraining. Third, the high cost of laboratory assays necessitates efficient computational methods that can infer previously unseen properties from existing experimental data, particularly by leveraging properties that have some degree of correlations. While some existing multimodal protein language models could infer new properties and handle diverse protein properties, they typically require fine-tuning or retraining for new laboratory conditions or assay formats.

In this work, we contribute to addressing these challenges by: (1) introducing a training strategy called *AB-context-aware* to force prompt-based models to condition their prediction on antibody sequence/property pairs examples provided in the context rather than relying uniquely on the sequences. This allows for directly accounting for batch effects that are represented in the context; (2) introduce *Context-aware Multi-Property Antibody Predictor or CA-MAP*, a multimodal architecture that features a specialized tokenization and embedding projection system for integrating text and protein language models. Unlike existing approaches such as TxGemma that rely on extensive fine-tuning of large foundation models, our approach enables prompting without dictionary merging across modalities, resulting in a compact model capable of in-context learning for multi-property prediction that can accept a variable number of antibody sequence/property values. Our approach relies on an orchestrating deep learning model which receives prompts uniquely composed of learnable tokens and various projectors and linear neural network layers to encode pLM, text and numerical value embeddings. This avoids the need of projecting the pLM embeddings into text space or learning amino acid sequences from scratch as text. The orchestrating model uses Mamba-based layers that use selective state spaces to perform content-based reasoning on long sequences more efficiently than standard transformer architectures. Antibody developability properties are not independent entities but rather interconnected characteristics that collectively determine therapeutic viability. For instance, hydrophobicity patterns may correlate with aggregation propensity, while charge distribution can influence both solubility and immunogenicity profiles. By incorporating multiple correlated properties in the prediction context, *CA-MAP* can capture these underlying relationships and make more informed predictions.

We show that our approach, trained from scratch with orders of magnitude fewer parameters, can achieve competitive performance compared to fine-tuned multimodal LLMs like TxGemma while offering superior adaptability to multi-property predictions and explicit batch effect handling capabilities.

## Results

### Dataset

These experiments used the extensive dataset generated by Bashour et al.^5^ The original data comprise of sequences primarily sourced from the Observed Antibody Space (OAS) database (1,738,091 sequences) and supplemented with experimentally-generated sequences (298,698 sequences). The dataset included human heavy chains (IgD, IgM, IgG, IgA, IgE) and light chains, as well as mouse heavy chains (IgM, IgG) and light chains. The authors calculated sequence-based and structure-based developability parameters for each chain of the antibody. In this work, we selected only heavy chains obtaining a total of 876,898 unique antibodies.

In this work, we selected a subset of heterogenous developability properties estimated some estimated with sequence-based predictors and some with structure-based predictors, specifically: Hydrophobicity (sequence-based hydrophobicity score with Peptides R package^29^), Solubility (sequence-based solubility index property with SoluProt v.1.0^30^), Stability (sequence-based aliphatic index computed with Peptides R package^29^), Immunogenicity (structure-based minimum rank property with netMHCIIpan version 4.0^31^), positive and negative charge heterogeneity (structure-based scores with therapeutic antibody profiler^6^). To avoid weighting properties differently throughout the experiments, they were normalized using fixed values such that even when different batch effects are simulated, they will still be between 0 and 1.

To avoid any chance of data leakage between training and testing, the sequences in the dataset were first split into three sets, setA (70% of total sequences), setB (15% of total sequences) and setC (15% of total sequences). All the sequences in the sets are unique. Antibody sequences and properties were sampled from each set independently to generate prompts and expected answers. The sampling procedure was entirely within set, such that prompts could not contain sequences from two different sets.

Each prompt contained two sections: *context* and *query*. The context section contains a variable number of antibodies sequences each one with a variable number of properties. Each property contains a name and a value. Example:

<antibody>

<seq>QVELVESGGGLVQPGGS…</seq>

<property name=”Hydrophobicity”>0.5</property>

<property name=”Stability”>0.1</property> [..other properties..]

</antibody>

<antibody>

<seq>EVQLVESGGGLVKPGGSLKLSCAASGF…</seq>

<property name=”Stability”>0.1</property>

</antibody>

[..other antibodies..]

The query section contains a sequence and the name of the property to predict. Example:

<query-antibody>

<seq>EVQLVESGGGLVQPGCAASGF…</seq>

<property name=”Stability” />

</query-antibody>

In the experiments, setA was used as training set, setB as validation set, and setC as test set. The antibodies used as context in test prompts all come from the test set to avoid any chance of data leakage. In experiments simulating batch effects, a different batch effect was introduced for each property type in each prompt.

### AB-context-aware Training

*AB-context-aware* promotes prompt-based models to condition their predictions on antibody sequence/property pair examples provided in the context, rather than relying solely on sequence information. This approach enables direct incorporation of batch effects present within the contextual examples. In these experiments, we compare the effect of AB-context-aware training with different degrees of batch effects on Tx-Gemma to predict Hydrophobicity with 10 to 15 antibodies and Hydrophobicity properties as context.

Using the 876,898 unique antibodies and training, validation and test sets described in the dataset section. The following number of unique prompts and expected answers were generated: 70,156 for training, 15,043 for validation, and 15,043 for testing. All prompts and expected answers uniquely referred to a single property: hydrophobicity.

In Fig. 1, we compare Tx-Gemma fine-tuned using standard fine-tuning, i.e. the unmodified prompts and expected answer vs. the same model fine-tuned with prompts and modified answers using the *AB-context-aware* strategy. We test the two Tx-Gemma fine-tuned models under three different conditions: without any batch effect, with an additive batch effect ranging from 0 to 0.1 (sampled from a uniform distribution), and an additive batch effect ranging from 0 to 0.3 (sampled from a uniform distribution).

**Fig. 1:**
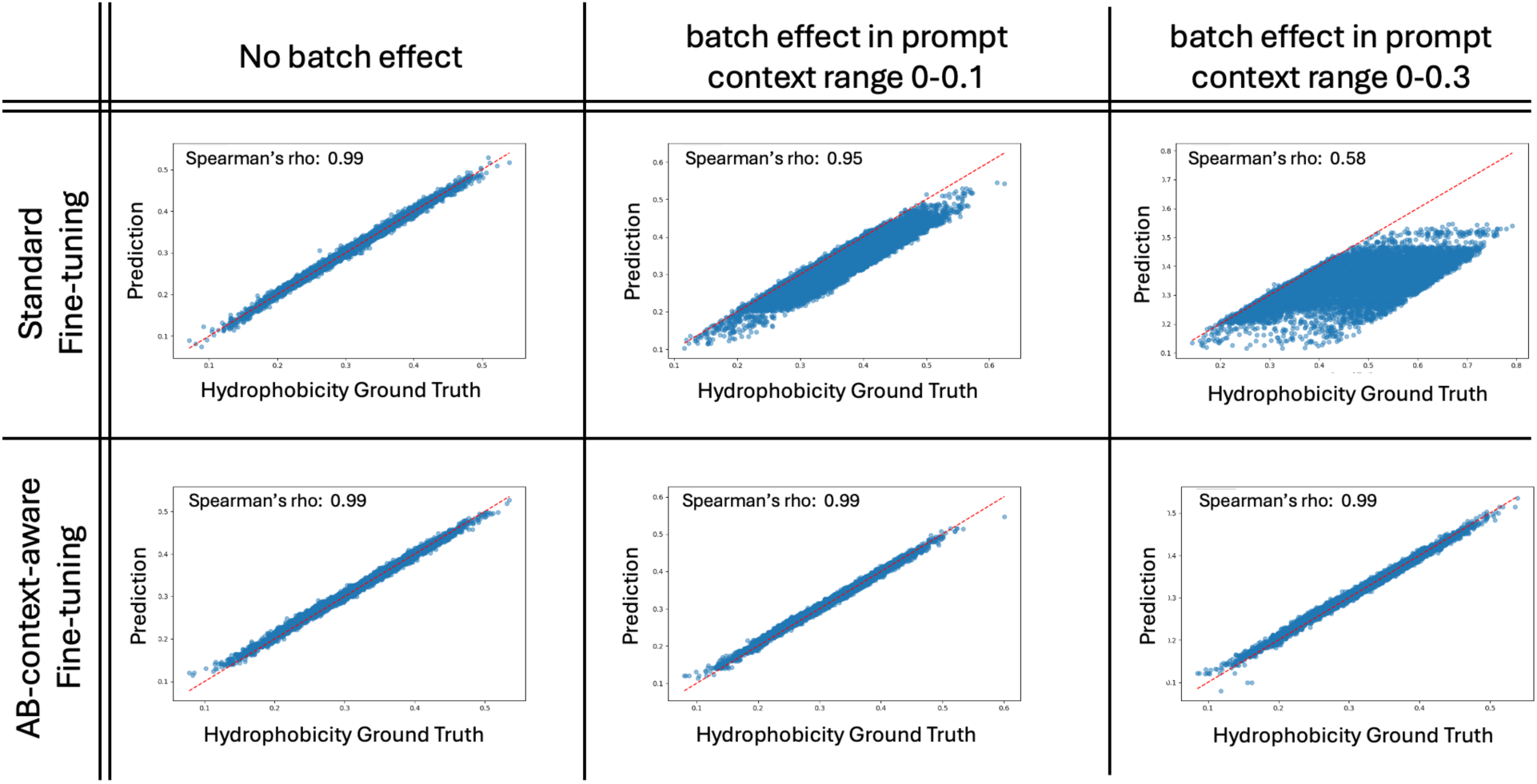
Comparison of Tx-Gemma with standard fine-tuning vs. AB-context-aware. Without the AB-context-aware fine tuning the model cannot infer the batch effect from the context.

Under ideal conditions without batch effects, both standard fine-tuning and AB-context-aware approaches achieved comparable performance (Spearman’s ρ ~ 0.99). However, as batch effect magnitude increased, the performance gap became pronounced.

With moderate batch effects (0-0.1 range), the standard fine-tuned TxGemma model showed substantial performance degradation (Spearman’s ρ = 0.95), while the AB-context-aware model maintained the same high predictive accuracy. This difference became even more dramatic under strong batch effects (0-0.3 range), where standard fine-tuning performance collapsed (Spearman’s ρ = 0.58) while AB-context-aware training preserved robust predictions.

In these experiments, Tx-Gemma produce parsable numerical answers in ~100% of cases when AB-context-aware fine tuning was used. While with standard fine-tuning it was ~100%, ~98%, ~88% for no bias, batch effect 0-0.1 and batch effect 0-0.3 respectively.

### CA-MAP and Multi-property prediction

In these experiments, we compare and contrast models trained to predict multiple properties. In Table 1 (top), we evaluate the performance of a fine-tuned Tx-Gemma model on predicting 4 developability properties (Hydrophobicity, NegCh heterogeneity, Solubility, Stability). The training set consisted of prompts generated using antibodies for setA, validation set with antibodies from setB and testing with antibodies from setC. Each prompt contained as context 10 to 15 antibody sequences with Hydrophobicity, NegCh heterogeneity, Solubility, Stability as properties. The query section contained a single sequence and property. This results in the following number of unique prompts and expected answers were generated: 70,156 for training, 15,043 for validation, and ~15,000 prompts for testing. Testing prompts are repeated 4 times, one per property queried. This is done to increase the number property queried given that Tx-Gemma struggles to output numerical properties given the prompts. In fact, we find that the maximum number of numerical results generated as output per property is 2.4% for Solubility. Solubility and Hydrophobicity are the only two properties where the model obtained statistically significant results, with ρ=0.1-0.18 for Solubility and ρ=0.74-0.79 for Hydrophobicity.

**Table 1:** Multi properties models comparison (Tx-Gemma and CA-MAP) and CA-MAP ablation study. Using multiple properties largely fails with Tx-Gemma. CA-MAP successfully predicts properties with or without bias. In CA-MAP ablation study, only the ESM2-embeddings are used to predict properties after tuning on the context (without the CA-MAP architecture). *** p<0.001; **p<0.01; *p<0.05; n.s. not significant.

|  | Tx-Gemma-multi-property<br>(AB-context aware) without<br>bias at inference time |  | Tx-Gemma-multi-property<br>(AB-context aware) with 0 to<br>0.3 bias at inference time |  |
| --- | --- | --- | --- | --- |
|  | *Spearman rho* | % predicted<br>(*n*) | *Spearman rho* | % predicted<br>(*n*) |
| **Hydrophobicity** | 0.74*** | 1.62% (224) | 0.79*** | 1.68% (253) |
| **NegCh**<br>**heterogeneity** | -0.3 (N.S.) | 0.11% (17) | -0.2 (N.S.) | 0.06% (10) |
| **Solubility** | 0.1* | 2.4% (374) | 0.18*** | 2.62% (394) |
| **Stability** | N/A | 0 | N/A | 0 |

Table 1: Multi properties models comparison (Tx-Gemma and CA-MAP) and CA-MAP ablation study.
|  | CA-MAP without bias at<br>inference time |  | CA-MAP with 0 to 0.3 bias at<br>inference time |  | ESM2-embedding only<br>(ablation experiment) |  |
| --- | --- | --- | --- | --- | --- | --- |
|  | *Spearman rho* | % predicted (*n*) | *Spearman rho* | % predicted<br>(*n*) | *Spearman rho* | % predicted (*n*) |
| **Hydrophobicity** | 0.95*** | 100% (3757) | 0.98*** | 100% (3757) | 0.42*** | 100% (3757) |
| **NegCh**<br>**heterogeneity** | 0.66*** | 100% (3683) | 0.84*** | 100% (3683) | 0.14*** | 100% (3683) |
| **Solubility** | 0.81*** | 100% (3758) | 0.83*** | 100% (3758) | 0.49*** | 100% (3758) |
| **Stability** | 0.97*** | 100% (3828) | 0.97*** | 100% (3828) | 0.47*** | 100% (3828) |

In Table 1 (bottom), we evaluate the performance of our CA-MAP model on predicting the same 4 developability properties. Training, validation and test sets were the same as the one used for Tx-Gemma. The only difference is that the test prompts did not need to be repeated 4 times, one per property queried, given that the model used ensured a numerical output by design. We find that all properties are predicted with statistically significant results. Hydrophobicity results into ρ=0.95-0.98, NegCh heterogeneity into ρ=0.66-0.84, Solubility into ρ=0.81-0.83, and Stability ρ=0.97. In this experiments, CA-MAP significantly outperforms the fine-tuned TxGemma for multi-property.

Given that CA-MAP encodes antibody sequences using ESM-2 embeddings^14^ as pLM, we also compare CA-MAP with prediction obtained using ESM-2 embeddings and a linear model fine-tuned on the properties used in each context used in the prompts. This acts as a naive baseline and allows to quantify the performance difference brought by the CA-MAP architecture. CA-MAP improvements over the naïve ESM-2 fine tuning on the properties used in the context ranged from ρ=0.34 to ρ=0.7.

CA-MAP including pre-trained ESM-2 and BERT sentence embedding components has a total of 125 million parameters. However, the vast majority of the parameters are frozen, resulting in ~182,000 trainable parameters. The Tx-Gemma model is the 9B version with an effective number of 5 billion parameters. All experiments with TxGemma in this work used Low-Rank Adaptation (LoRA) to reduce the number of trainable parameters to 40 million. Using a machine with a single NVIDIA L40S GPU with 46GB of memory, the running time at inference for the fine-tuned TxGemma is 22s per iteration (where each iteration has 12 prompts, equivalent to 1.83 s/prompt), while CA-MAP is 1.16s per iteration (where each iteration 128 prompts, equivalent to 0.009 s/prompt, however this figure does not include the time for creating sequence and text embeddings, which were precomputed).

### CA-MAP and Properties Unseen During Training

The developability characteristics of antibodies represent interconnected molecular features rather than isolated parameters, collectively influencing therapeutic potential. Molecular interactions exemplify these relationships. CA-MAP has the potential to incorporate multiple correlated properties within their predictive context, they can exploit these inherent molecular relationships to generate more accurate predictions for unknown properties.

The value of leveraging property correlations becomes most apparent when evaluating model performance on properties excluded from the training phase. As shown in Table 2, we assessed prediction accuracy for immunogenicity and positive charge (PosCh) heterogeneity under two distinct contextual scenarios: utilizing only these two target properties as contextual information versus employing the complete six-property dataset. The substantial performance enhancement—with Spearman’s ρ increasing from 0.25 to 0.73 for immunogenicity and from 0.08 to 0.73 for PosCh heterogeneity—when incorporating additional correlated properties indicates that CA-MAP can leverage molecular property interdependencies for enhanced predictive accuracy.

**Table 2:** We test the effect of adding more properties in the context by comparing the predictive ability of CA-MAP model on two properties that were not used in the training dataset: Immunogenicity and Positive Charge Heterogeneity. *** p<0.001; **p<0.01; *p<0.05; n.s. not significant.

|  | Full Context<br>(6 properties) |  | Immunogenicity and PosCh heterogeneity<br>as Context |  |
| --- | --- | --- | --- | --- |
|  | Spearman rho | % predicted (n) | Spearman rho | % predicted (n) |
| Immunogenicity | 0.73*** | 100% (749) | 0.25*** | 100% (710) |
| PosCh heterogeneity | 0.73*** | 100% (7597) | 0.08*** | 100% (7418) |

Both immunogenicity and PosCh heterogeneity were not used in the training or validation set, so all the model could use for prediction was the context, and what it has learned on the other four properties (in the experiment with full context). All experiments in Table 2 used the simulated 0 to 0.3 bias at inference time and antibodies from setC (the test set).

CA-MAP relies on ESM-2 embeddings for representing sequences. In Fig. 2, we evaluate the performance comparison between CA-MAP and an ESM-2-based linear model as a function of an increased number of antibodies used for context (or training in the case of ESM-2-based linear model). We picked PosCh Heterogeneity as target as it was not used in the training or validation set for CA-MAP training and there are enough sample in the training set to allow for an increase of antibody context without having to recycle them across prompts. using CA-MAP also included in the context other 5 properties (Hydrophobicity, NegCh heterogeneity, Solubility, Stability, Immunogenicity). CA-MAP has a clear performance advantage over ESM-2, this is likely because of its ability of leveraging correlation between properties. In both cases, performance improvements tend to flatten at the scale of 100 antibodies.

**Fig. 2:**
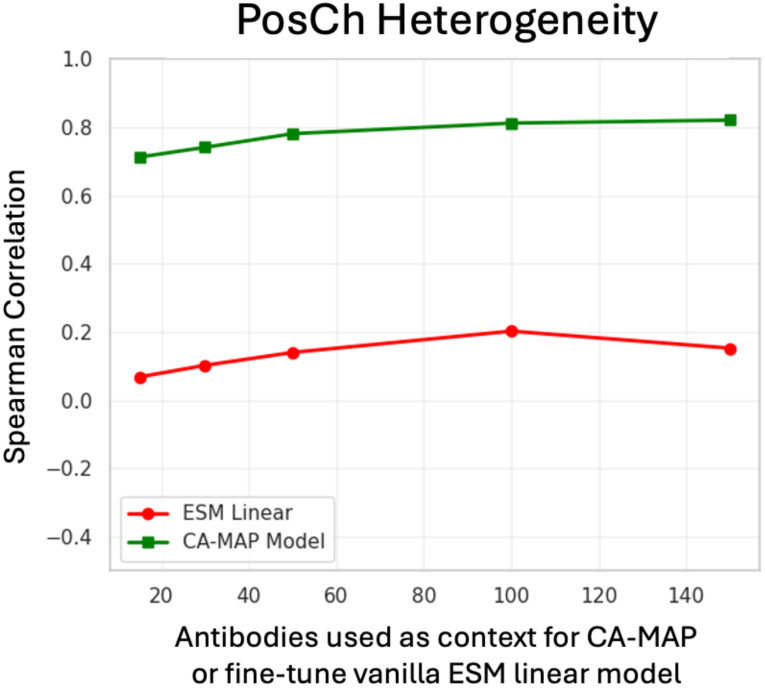
CA-MAP and linear model based on ESM-2 embeddings performance as a function of antibodies used as context (CA-MAP) or used to fine-tune ESM-based linear model.

## Discussion

The AB-context-aware training strategy addresses a critical limitation in current machine learning approaches to antibody property prediction: the tendency of models to ignore contextual information and rely exclusively on sequence data. Our results demonstrate this. Under ideal conditions without batch effects, both standard and context-aware fine-tuning achieved comparable performance (Spearman’s ρ ≈ 0.99). However, as batch effect magnitude increased, the performance gap became decisive. With moderate batch effects (0-0.1 range), standard fine-tuning degraded to ρ = 0.95 while context-aware training maintained ρ > 0.99. Most strikingly, under strong batch effects (0-0.3 range), standard fine-tuning collapsed to ρ = 0.58 while context-aware maintained ρ ≈ 0.99. This mechanism works by introducing latent transformations during training that create explicit dependencies between context and predictions, forcing the model to internalize how batch effects manifest in contextual examples. This training constraint transfers to inference, enabling the model to account for batch effects through in-context examples without requiring explicit statistical methods.

The transition from single-property to multi-property prediction reveals that our proposed architecture, CA-MAP, achieves statistically significant predictions across all four tested properties, whereas TxGemma produced parsable numerical outputs for only 2 of 4 properties with lower correlation in all cases. This performance gap reflects a fundamental architectural principle: maintaining separate embedding pathways for different modalities could preserve domain-specific knowledge. Rather than projecting protein embeddings into text space or vice versa, CA-MAP treats sequences, property names, and numerical values distinctly through learnable projectors, enabling the Mamba orchestrator to compose information across modalities. CA-MAP’s computational efficiency further enhances practical applicability. With smaller number of trainable parameters and significantly faster inference, the model enables iterative in-the-loop workflows and deployment on modest hardware.

Compared to text-based multimodal large language models such as TxGemma, CA-MAP maintains distinct modality pathways that avoid information loss from cross-modal projections. It eliminates the need for extensive fine-tuning by enabling context-based adaptation at inference time, uses orders of magnitude fewer trainable parameters, and guarantees numerical output without requiring parsing of model outputs. When compared to fine-tuned linear models applied directly to embeddings, CA-MAP captures complex relationships through Mamba’s selective state spaces, leverages multi-property correlations in the context, and generalizes better to unseen properties and batch conditions. Compared to standard protein language models applied alone, CA-MAP incorporates property semantics through BERT projections, enables batch effect handling through context provision, and makes multi-property predictions possible from a single model rather than requiring separate models per property.

A particularly significant finding is that antibody developability properties share exploitable correlations. When predicting unseen properties (immunogenicity and positive charge heterogeneity), incorporating additional correlated properties improved performance dramatically. These improvements reflect the physiochemical interconnection of developability properties—hydrophobicity determines aggregation propensity, charge distribution influences solubility and immunogenicity, and backbone conformation affects both stability and immunogenicity. This capability has substantial practical value. If costly assays can be predicted from cheaper ones plus sequence information, a mutation campaign with thousands of antibodies could achieve substantial cost reduction and accelerated decision-making.

The practical applicability of CA-MAP extends to integration within agentic and generative design frameworks. The model’s efficiency and guaranteed numerical output make it suitable for rapid inference in iterative loops where computational predictions guide experimental prioritization. Unlike heavy foundation models, CA-MAP enables few-shot adaptation to new laboratory conditions by simply including batch examples in the context, eliminating the need for retraining. This flexibility combined with sub-second inference times opens possibilities for real-time decision-making in high-throughput screening campaigns, where thousands of candidate predictions must be generated and ranked within minutes rather than hours.

Understanding the mechanistic basis of property correlations leveraged by CA-MAP remains an important open question. While the model demonstrates strong predictive performance, the underlying molecular relationships driving these correlations warrant deeper investigation through attention visualization, ablation studies on individual projector layers, and systematic analysis of learned representations. Furthermore, the generalizability of the context-aware approach to other protein engineering domains beyond antibodies—such as enzyme engineering, scaffolding proteins, or cell therapies—represents a natural extension warranting exploration. The framework’s ability to handle missing data and variable context sizes also suggests applicability to real-world laboratory settings where incomplete assay panels are commonplace, though this requires more experimental validation.

However, important limitations must be acknowledged. Our dataset comprises computationally predicted rather than experimentally measured developability properties, still all of the results reported can also be interpreted in relative terms, rather than absolute performance, where the relative changes between experimental setups, baseline and models are likely to hold regardless of the dataset type. Additionally, the model’s predictions depend critically on context quality and representativeness. Our work was limited to six biochemical properties; generalization to binding affinity, specificity, or off-target binding remains unexplored. Experimental validation with additional laboratory data across diverse conditions and facilities is essential before broad adoption in development. Despite these limitations, the framework suggests practical implications for antibody development: property prediction cascades where cheap assays inform selection of expensive ones, rapid adaptation to laboratory-specific conditions through few-shot context examples, and integration into generative design frameworks.

## Methods

### AB-context-aware Training

A promptable model like a multi-modal LLM would typically be trained/fine-tuned by presenting multiple examples of prompts/answers and penalizing any incorrect answer with some type of loss. However, if the model is trained using a dataset coming from the same source, it can easily lead to shortcut learning as the model could simply ignore the context and just use the query antibody sequence for its prediction. In AB-context-aware, we introduce a hidden latent variable in the properties/features of the antibody used as context and the answer. This latent variable changes for each prompt used in the training set. This ensures that the only way the model can predict the properties/features correctly is if it has considered the context. Fig 3 shows the intuition of the method. An example of this latent variable could be an additive constant to the properties of the antibodies, but it could also be a nonlinear transformation parametrized by the latent variable, i.e. new_property = f(original_property; latent_variable).

**Fig. 3.**
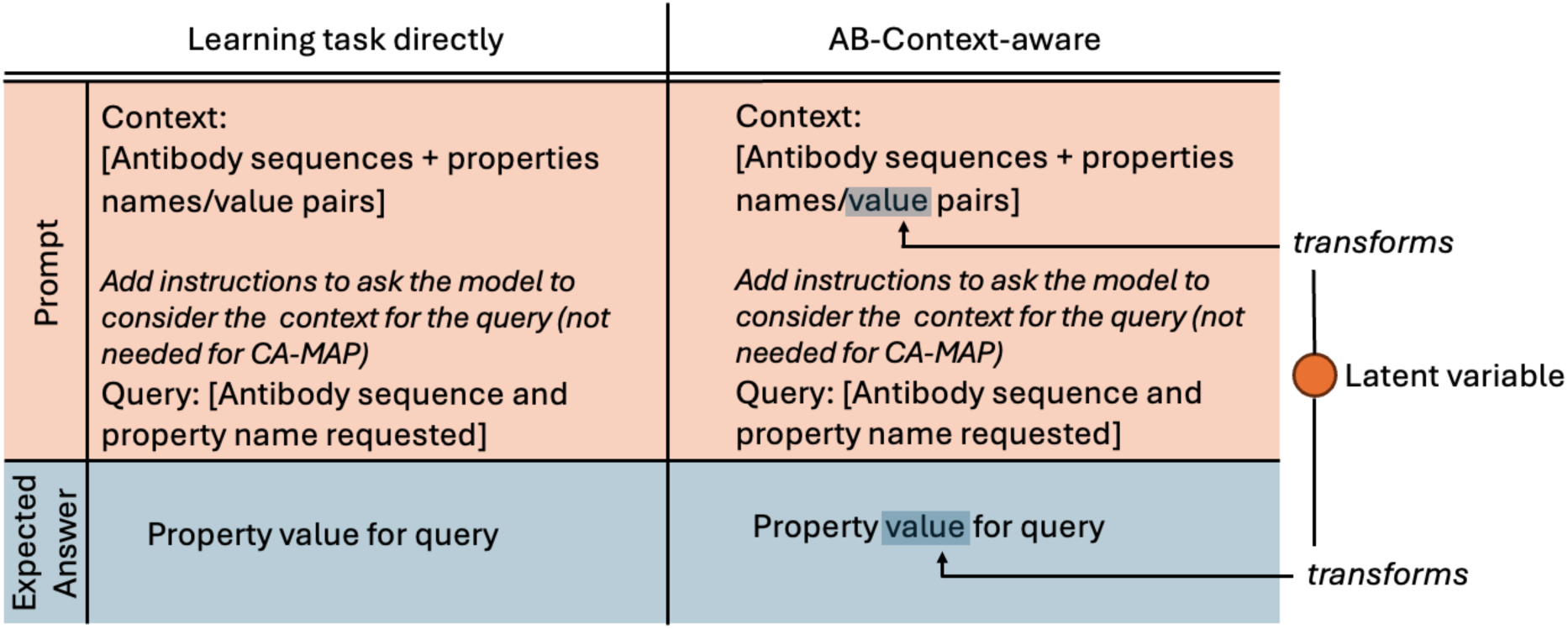
High-level prompts and expected answers. Right column shows the standard fine tuning process where the task is learned directly; left column, shows the intuition behind AB-context-aware training strategy where a model is forced to consider the context to obtain a correct property value instead of relying uniquely on the query antibody sequence.

To avoid introducing new learning shortcuts on the latent variable, it is important to have a latent variable that is randomly picked each time a training prompt is generated.

Formally, we seek a model that predicts property values y given query q (containing amino acid sequence) and context ctx (containing example sequences/properties):

### P(y | q, ctx)

However, models may ignore context through shortcut learning, effectively learning: P(y | q, ctx) ≈ P(y | q)

This occurs when training datasets from the same experimental campaign make y approximately conditionally independent of ctx given q.

This is problematic because: (1) we want models to adapt to new data in ctx, and (2) this prevents accounting for batch effects.

To force context dependence, we introduce transformation f_tr(x, l) where l is a random latent variable sampled during training. By design, f_tr ensures:

P(f_tr(y, l) | q, f_tr(ctx, l)) ≠ P(f_tr(y, l) | q)

This is achieved because f_tr creates dependence between transformed outputs and context: P(f_tr(y, l) | f_tr(ctx, l)) ≠ P(f_tr(y, l))

This can be implemented with the following python-like pseudocode

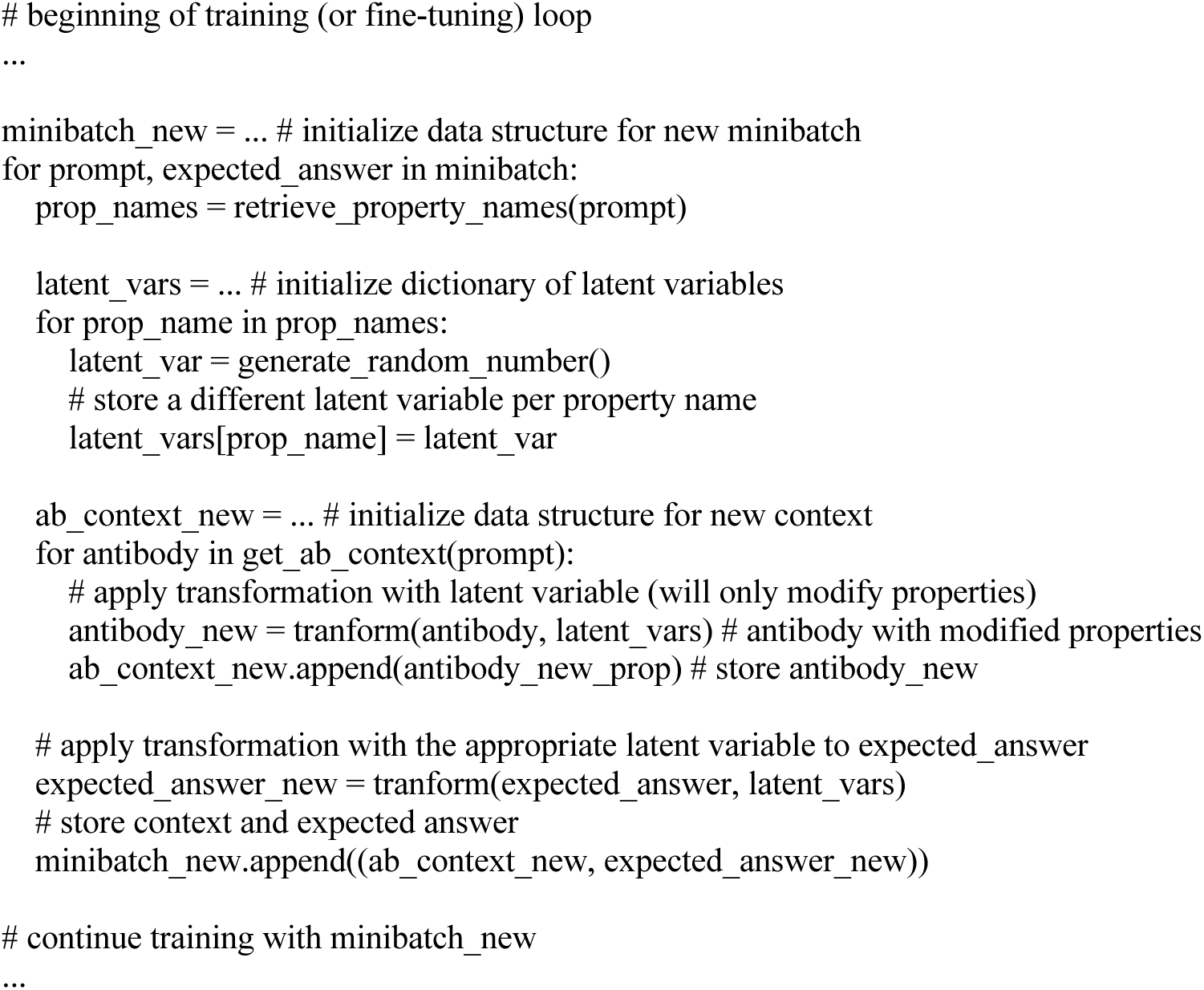

### *CA-MAP* Description

CA-MAP (Context-aware Multi-Property Antibody Predictor) aims to be a compact multimodal architecture designed to integrate protein language models (pLMs) with natural language embeddings for multi-property antibody prediction with in-context learning capabilities. Unlike conventional approaches that project embeddings across modalities, CA-MAP maintains separate embedding pathways for distinct data types including protein sequences, text, and numerical values. This design enables the orchestrating Mamba-based^32^ model to compose information across modalities without loss of domain-specific knowledge. The key innovation in CA-MAP’s design is its specialized tokenization scheme that converts the prompt into a structured sequence rather than natural language text and the explicit use of context and query sections. This approach eliminates the need for dictionary merging across modalities, substantially reducing memory requirements and computational overhead. The tokenization strategy accommodates variable numbers of antibodies, properties, and missing data within a single framework, and enables flexible querying of any property at inference time without model retraining.

As shown in Fig. 4(a), the prompt structure for CA-MAP is organized into two main sections. The context section contains multiple antibody examples, each with an amino acid sequence tag followed by multiple associated properties with their values. This design supports a variable number of antibodies and properties, enabling the model to handle incomplete data. The query section contains the target antibody sequence to predict for, along with the name of the property to predict, with the property value to be determined at inference time. Learned special tokens serve as delimiter vectors with learnable embeddings of dimensionality 32. These tokens, such as eos tokens, antibody tags, and query-antibody tags, signal structural transitions to the model, enabling it to understand the prompt hierarchy and process information appropriately.

**Fig. 4.**
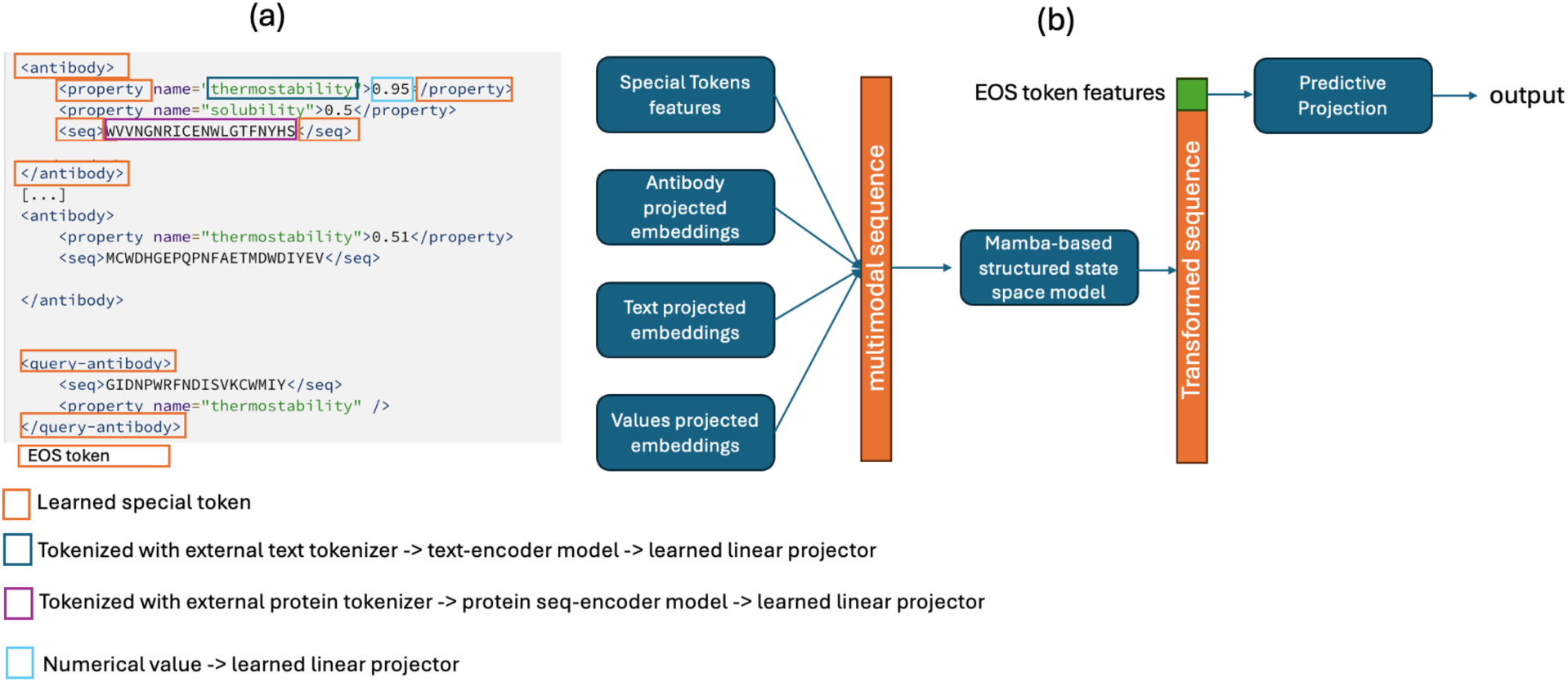
CA-MAP architecture diagrams. (a) Shows how the different components of the prompt are tokenized and encoded; (b) Overview of the model showing the 4 main streams of encoded features from: special tokens, antibody projected embeddings, text projected embeddings, and numerical values projected embeddings.

The architecture receiving this tokenized input consists of multiple parallel embedding pathways that feed into a Mamba-based orchestrator. The first pathway handles antibody sequences through a pre-trained protein language model (ESM-2^14^, pre trained checkpoint: esm2_t6_8M_UR50D) with frozen weights to preserve learned protein representations. The ESM-2 encoder produces a vector representation of the last hidden state for each residue, which are then averaged across all residues to produce constant-length embeddings regardless of sequence length. These embeddings are compressed through a learnable linear transformation to a target dimensionality of 32, allowing the model to adapt pre-trained representations to the specific task.

The text pathway processes property names through pre-trained Sentence Embeddings using Siamese BERT-Network^33^ (checkpoint: sentence-transformers/paraphrase-multilingual-MiniLM-L12-v2). With frozen weights to preserve semantic understanding, this encoder takes property names such as “Hydrophobicity” or “Stability” and produces semantic embeddings via mean pooling of token representations. Like the antibody pathway, these embeddings are then compressed through a learnable linear transformation to dimensionality 32, enabling the model to learn task-specific semantic relationships between properties.

The numerical values pathway represents property measurements. Single floating-point numerical values are expanded through a learnable linear transformation to dimensionality 32, allowing the model to learn appropriate scaling and embedding of numerical values. This pathway supports property values normalized to the range of 0 to 1. Finally, the remaining pathway comprises the learned special token embeddings themselves, which also have dimensionality 32. These token embeddings are trainable parameters initialized randomly and updated during training, marking structural elements and transitions throughout the prompt. The four parallel embedding pathways converge into the orchestrating model based on Mamba architecture, which represents a selective state space approach for efficient language modeling. This model is configured with 6 layers, 32 hidden states, a local convolutional width of 4, and a block expansion factor of 2. Importantly, the model is fully trained from scratch, unlike the frozen embedding layers, allowing it to learn how to compose information across the different modalities. The architecture, being based on Mamba offers several advantages over standard approaches: no positional encoding is required since Mamba’s selective state spaces encode position information implicitly through state transitions, eliminating the need for explicit positional embeddings. Additionally, it allows for efficient long-sequence modeling with linear-time complexity relative to sequence length, enabling the processing of long prompts with many context examples. Data flows through as concatenated sequences of special tokens (learned embeddings), antibody embeddings (from the ESM-2 projector), text embeddings (from the BERT projector), and value embeddings (from the numerical projector). The prompt processes through 6 sequential Mamba layers, with each layer performing selective state space transformations, content-based filtering and reasoning, and progressive refinement of representations. The outputs from these layers are contextualized representations for each input element.

The prediction head of CA-MAP consists of a final learnable linear transformation applied to the output from the Mamba layer associated with the eos token, producing a single numerical value representing the predicted property value. Building the prediction head on the last special token, rather than an initial one, is a requirement of state space models based on Mamba to enable the token to attend to all the previous ones. A critical advantage of CA-MAP is its multi-property capability achieved through this single unified prediction head design. Unlike conventional multi-task models that require task-specific prediction heads, CA-MAP uses a single head to achieve property-agnostic learning that works across different developability properties. This design enables inference-time flexibility, allowing the model to predict any property by simply specifying the property name in the query section. Moreover, the model can adapt to different batch conditions by including batch examples in the context without requiring retraining.

Each modality, sequence, text, and numerical, maintains a distinct embedding pathway rather than forcing information through cross-modality projections. This approach avoids problematic projections of protein embeddings into text space or vice versa, while preserving each modality’s pre-trained knowledge through frozen weights. The model is relatively compact with a total of 125 million parameters, with the vast majority of these coming from the frozen ESM-2 and BERT components. Only approximately 182,000 parameters are trainable during model training. This parameter efficiency achieves competitive performance with orders of magnitude fewer trainable parameters compared to foundation models even when fine-tuned with Low-Rank Adaptation (LoRA),^34^ allowing directly training from scratch without a pre-training step.

CA-MAP allows for variable numbers of antibodies and properties per context, supports missing data where properties can be omitted for specific antibodies. For multi-property prediction, the model is trained once on multiple properties, avoiding the need for property-specific fine-tuning. Because of this design, CA-MAP can predict properties not seen during training as long as it can leverage correlations with properties learned already and the ones provided in the context examples.

## Statistics and reproducibility

The study utilized Spearman’s correlation analysis, computed through scipy.stats library functions. No power analysis was performed to determine sample sizes in advance, and the analysis incorporated all available data without any exclusions. The study protocol did not implement randomization, and investigators conducted experiments and assessments without blinding to allocation.

## Data visualization

All figures were created using a combination of software tools including Python, matplotlib, seaborn, and PowerPoint.

## Data availability

Raw antibody sequence and property data is available at https://github.com/csi-greifflab/developability_profiling/blob/main/data/native/developability.csv and was created by the authors of Bashour et al. ^5^

Code is available at https://github.com/amazon-science/ca-map

## Supplementary Information

### CA-MAP training

CA-MAP was trained with a mean squared error loss for a maximum of 20 epochs and weights were selected based on the best loss on the validation set. The AdamW algorithm was used as optimizer with batch_size = 128, learning_rate = 0.001, and weight_decay=0.01.

Example loss curves are shown in Fig. S1

**Figure S1.**
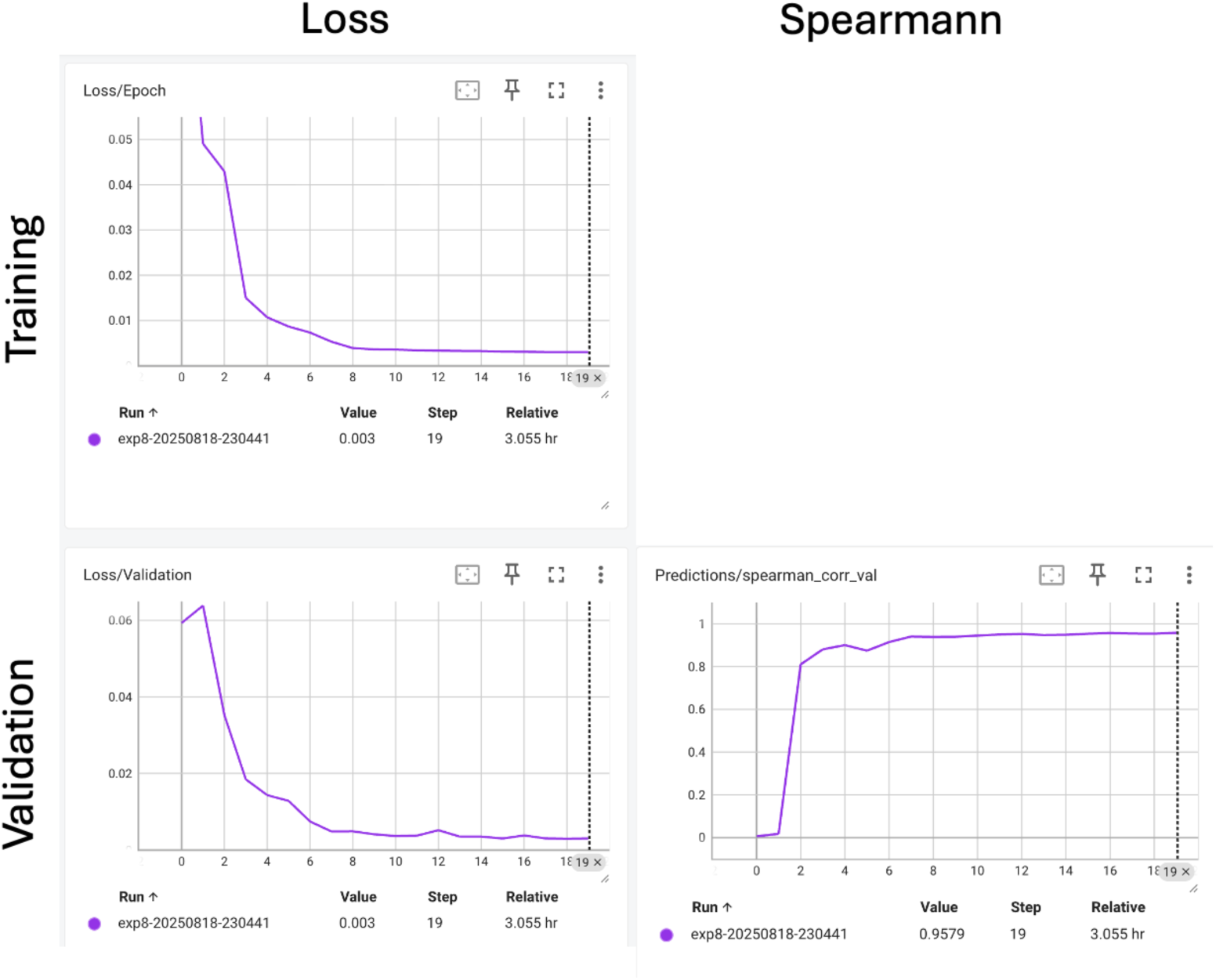
Example CA-MAP loss curves

### TX-Gemma

TxGemma was trained using LoRA using a rank of 12 targeting the following components of TxGemma “q_proj”, “o_proj”, “k_proj”, “v_proj”, “gate_proj”, “up_proj”, “down_proj”. The optimizer used was AdamW with a batch_size=4, gradient_accumulation_steps=4, warmup_steps=2, learning_rate=2e-4. The model was trained for over 1 epoch and weights were selected based on the best loss on the validation set. Preliminary experiments increasing the training time to up 4 epochs did not change the performance. Example loss curves are shown in Fig. S2.

**Figure S2.**
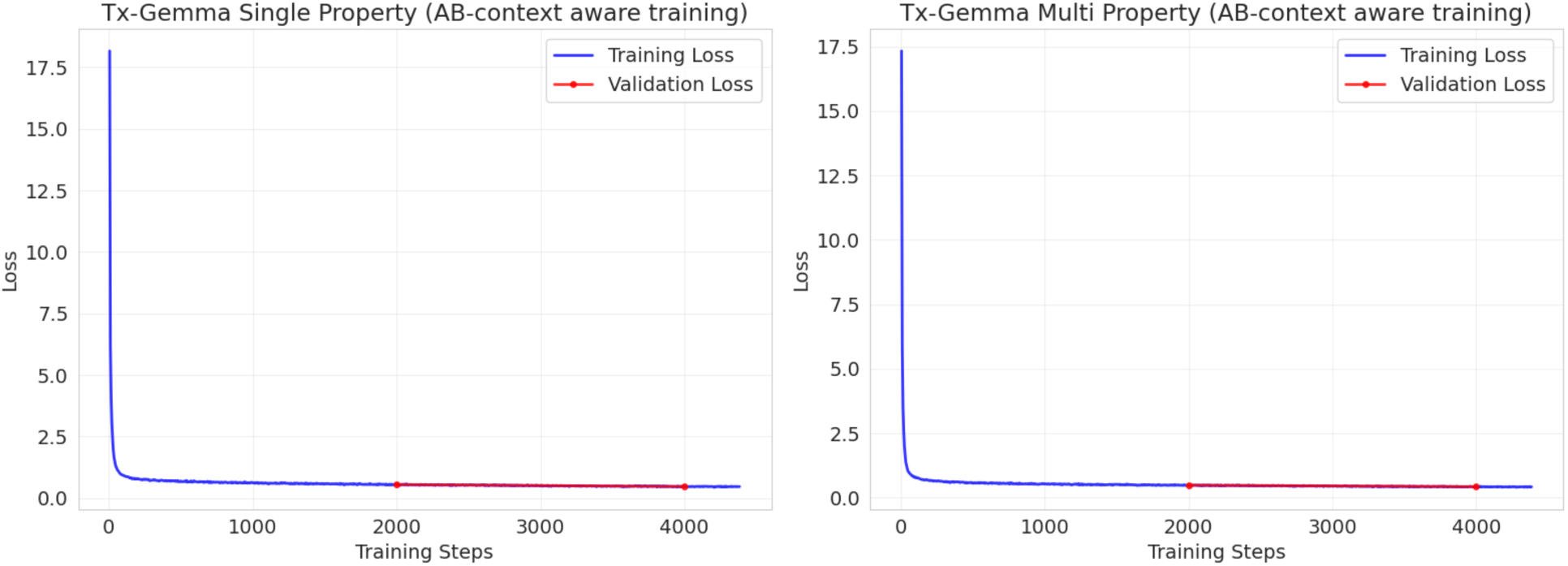
Example Tx-Gemma loss curves

### Property Normalization

To avoid weighting properties differently throughout the experiments, they were normalized using fixed values such that even when different batch effects are simulated, they will still be between 0 to 1. The following values were used for normalization:

Solubility: {min: 0, max: 1}

Immunogenicity: {min: 0, max: 2}

Stability: {min: 30, max: 100}

Hydrophobicity: {min: −0.2, max: 0.6}

PosCh heterogeneity: {min: 0, max: 10}

NegCh heterogeneity: {min: 0, max: 10}

